# Binomial models uncover biological variation during feature selection of droplet-based single-cell RNA sequencing

**DOI:** 10.1101/2021.07.11.451989

**Authors:** Breanne Sparta, Timothy Hamilton, Samuel D. Aragones, Eric J. Deeds

## Abstract

Single-cell RNA sequencing (scRNA-seq) aims to characterize how variation in gene expression is distributed across cells in tissues and organisms. Yet, effective comprehension of these extremely high-dimensional datasets remains a critical barrier to progress in biological research. In standard analyses of scRNA-seq data, feature selection steps aim to reduce the dimensionality of the data by focusing on a subset of genes that are the most biologically variable across a set of cells. Ideally, these features provide the genes that are the most informative for partitioning groups of transcriptionally distinct cells, each representing a different cell type or identity. In this work, we propose a simple feature selection model where a binomial sampling process for each mRNA species produces a null model of technical variation. To compare our model to existing methods, we use scRNA-seq data where cell identities have been established *a priori* for each cell, and characterize whether different feature sets retain biologically varying genes, distort neighborhood structures, and allow popular clustering algorithms to partition groups of cells into their established classes. We find that our model of biological variation, which we term “Differentially Distributed Genes” or DDGs, outperforms existing methods, and enables dimensionality reduction without loss of critical structure within the data set.

## Introduction

Single-cell RNA sequencing has advanced the resolution at which variation in gene expression can be observed [Kalisky 2018]. Recent improvements in scRNA-seq technology have enabled the measurement of tens of thousands of genes across hundreds of thousands to millions of cells [Svensson 2018]. Yet, interpreting transcriptional variation in these extremely high-dimensional datasets remains a challenge at the forefront of biological research [Kiselev 2019].

In addition to the high levels of biological variation observed in scRNA-seq data, the sparsity of the data can pose challenges during downstream analysis procedures [Lun 2016]. There exist very small quantities of mRNA inside of single cells, and even with advanced microfluidic technologies, the capture probability for any given mRNA is low [Shapiro 2013]. As a result, a typical scRNA-seq experiment can generate a gene-by-cell expression matrix where ∼95% of the entries are zeros. The observance of this large fraction of zeros, colloquially termed “drop-outs”, has left an impression that scRNA-seq data is zero inflated [Finak 2015, Pierson 2015, Liu 2016, Lin 2017, Risso 2018]. However, there is a growing body of empirical and theoretical work that demonstrates that, given the distributions of gene-count data, we do not observe more zeros in scRNA-seq data than would be expected based on a sampling process where the probability of capturing any given mRNA molecule is low [Townes 2019, Svensson 2020, Sarkar 2021].

A major contributor to this observation is the fact that droplet-based approaches reduce technical error in scRNA-seq [Macosko 2015, Klein 2015, Zheng 2017, Svensson 2017]. In these experiments, individual mRNA’s from lysed cells are bound to Unique Molecular Identifiers, or UMI’s, prior to PCR-based amplification, thus overcoming amplification bias when measuring mRNA abundance [Islam 2014]. Despite these advances, scRNA-seq approaches still entail a low capture probability for individual mRNA molecules, and the resulting sparseness of this high-dimensional data pose challenges for the analysis of biological variation [Kiselev 2019]. Standard approaches to scRNA-seq data analysis often employ a feature selection step where observed variation in gene expression values is compared to a model of measurement error [Brennecke 2013, Andrews 2019]. The goal of this feature selection step is to identify genes whose expression levels are varying due to meaningful biological differences in the population, rather than due to just technical noise. This step often significantly reduces the dimensionality of scRNA-seq data to only contain biologically varying genes prior to partitioning cells into transcriptionally similar groups.

While many null models of biological variation have been proposed, which feature selection method is most appropriate remains unresolved [Duo 2018, Andrews 2019, Kim 2020, Su 2021]. Software that is commonly referred to as the “best practice” approach to scRNA-seq analysis, Seurat and Scanpy, use a feature selection procedure that identifies “Highly Variable Genes” or HVGs [Luecken 2019, Butler 2018, Wolf 2018]. In this procedure, the raw UMI counts in the data are first transformed with a counts-per-million (CPM) normalization and then with a log+1 transformation. The motivating idea of these transformations are to normalize for cell-size factors and stabilize the variance for genes whose averages are orders of magnitudes different, respectively. However, both transformations have the undesirable effect of increasing the distance between 0 and non-zero values within a gene’s distribution [Lun 2016, 2018, Townes 2019]. As a result, these transformations artificially increase the variance of genes, and genes with mean counts that are closer to zero are disproportionality inflated [Townes 2019]. Further, CPM normalization aims to adjust counts for potential read depth-differences that occur from differences in cell size. However, in droplet-based approaches, the assumption that cell-size can affect read-depth lacks empirical support. As a result, both of these transformations can introduce bias into variance estimates, and potentially skew downstream cell-clustering and analysis results. After performing these transformations, a generalized linear model is used to fit the relationship between the mean and variance across all the genes. Genes with variances that are considerably higher than would be predicted by this model on the basis of their average expression are considered “highly variable.” These models generally do not provide a natural cutoff for how many such genes to include, however, and as such the number of HVGs is an essentially arbitrary choice left to the user. Most practitioners select between 1000 and 5000 HVGs to focus on for further analysis [Luecken 2019].

While the approach sketched above is common in the literature, there is no theoretical basis to expect that the HVG approach captures a set of genes that varies in the population due to biological factors. Also, as mentioned above, normalization, log transformation and the choice of the number of HVGs to consider all have the potential to introduce biases into downstream analysis. As a result, there is an ongoing effort in the field to improve feature selection in scRNA-seq. One common alternative approach is to model the abundance of zeros, or ‘dropouts’, in gene distributions and to identify features that have more zeros than expected. For instance, a recent software package developed by Andrews and Hemberg offers three different null models that develop an expectation for the relationship between the ‘dropout’ rate and the mean expression level of each gene: M3Drop, NBDrop, and NBDisp. In M3Drop, a dropout rate parameter is fit to the whole transcriptome, and a Michaelis-Menten-style hyberbolic function is used to identify outliers where the gene-specific dropout rate exceeds the population expectation [Andrews 2019]. NBDrop uses a negative binomial distribution to model the fraction of zeros per gene as a function of the mean gene count that is adjusted for the sequencing efficiency, or total number of counts, for each cell. NBDisp uses the same negative binomial model as NBDrop, but is similar to the HVG method in that it uses linear regression to model the relationship between the mean and the estimated dispersion to identify a set of overly-dispersed genes. In another method similar to the NBDrop approach, Townes et al. identified a set of genes with a greater fraction of zeros than is expected, using a multinomial model where the probability of a mRNA being captured depends on its relative abundance within each cell [Townes 2019].

In each of these previous studies, feature selection models are evaluated based on clustering performance using biological data with “ground truth” cell identities, or by their ability to recover differentially expressed genes that have been identified using bulk RNA-seq methods. Using these approaches, it has been demonstrated that the HVG method only marginally improves performance compared to a randomized feature selection control [Andrews 2019]. Yet, the HVG method remains extremely popular [Luecken 2019, Park 2018, Fincher 2018, Gerber 2018, Siebert 2019, Xi 2020, Diaz 2020, Collin 2021, Fawkner-Corbett 2021, Li 2021]. In addition to uncertainty about the ability of various methods to identify an informative set of biologically varying genes, the HVG, NBDisp, and Townes method all entail arbitrary decisions about the number of features to use in the downstream clustering procedure. Because the choice of feature selection models can significantly alter a study’s results, there is a need to further develop statistically-grounded feature selection approaches, and to subject these methods to rigorous tests that evaluate their capacity to identify genes that exhibit *bona fide* biological variation within a population.

In this work, we proposed a simple binomial model that can be used to identify differentially distributed genes, or DDGs. In this model, each mRNA has the same probability of being captured during the initial phase of the scRNA-seq experiment, which results in a binomial sampling process of the mRNA molecules within the cell. We then consider the simple null model where there is no biological variation in a gene’s expression across the population, and then calculate the probability that purely technical variation due to the mRNA capture process could generate the observed expression patterns. While similar in spirit to the multinomial model developed by Townes et al., our inclusion of a specific null model allows us to calculate a *p*-value that represents the chance that the observed variation in a gene’s expression could be explained purely by technical noise. As a result, our approach allows researchers to define a standard False Discovery Rate (FDR) that sets the number of false positives they are willing to accept, rather than an arbitrary cutoff in the number of genes to consider for further analysis.

We validated this model using a standard synthetic RNA spike-in scRNA-seq data set, where all of the variation should arise from the sampling process. We found that the vast majority of these genes were consistent with the null model, as expected. We further tested our approach by clustering cells from a study of FACs-sorted lymphocytes, where cell identity markers are obtained prior to scRNA-seq [Zheng 2017]. We compared our model to a number of existing feature selection approaches, and found that DDGs better retain the structure of variation among established cell identities and more accurately identified genes that are differentially expressed between cell types. DDGs also provide a feature set that better partitions groups of cells in accordance with their established cell identity labels. We found that, while feature selection only marginally improves cell clustering performance compared to the full feature set, the DDG approach enables dimensionality reduction without loss of critical structure within the data set. Overall, our findings suggest that DDGs can more accurately and comprehensively identify genes whose expression patters demonstrate true biological variation compared to existing methods.

## Results

### Characterizing patterns of gene expression variation across tissues and multicellular organisms

Single-cell RNA sequencing technology has promised to map functional diversity by quantifying global gene expression patterns at the resolution of individual cells. This enhanced resolution has the potential to revolutionize our understanding of how gene expression and cellular identities are related. Yet elucidating organizational principals of physiological diversity in multicellular tissues remains an ongoing challenge. In scRNA-seq experiments, the dominant approach to quantifying transcriptional variation is to identify genes that are differentially expressed across groups of transcriptionally similar cells. Here, transcriptionally similar cells are identified by computing the distance between cells within a coordinate space that is defined by gene expression counts, and clustering cells that occupy dense regions in gene expression space [Luecken 2019, Krzak 2019, Qi 2020]. To gain intuition about the structure of transcriptional variation, we took the inverse approach, and sought to characterize groups of genes that are expressed across similar sets of cells. To do this, we transposed the cell by gene expression matrix, and defined distance between genes as distance in expression values across the set of cells. To identify and compare communities of genes that are expressed in similar patterns across complex tissues, we performed gene-clustering multiple times across a diverse set of scRNA-seq data sets.

Louvain clustering was performed on the gene features of scRNA-seq data collected from human lymphocytes, mouse bladder, mouse kidney, hydra, or planaria (Figure 1, S1) [Blondel 2008, Zheng 2017, Li 2021, Park 2018, Siebert 2019, Fincher 2018]. Louvain clustering was performed on each sample independently, and across these samples we observed three general expression patterns: 1) A group of genes that are nearly ubiquitously expressed across all cells. These genes are comprised largely of ribosomal proteins, and genes involved in energy production, such as the ATP-synthase. 2) Sparsely expressed genes that are found in few, non-overlapping sets of cells, with no apparent structure in the expression pattern. 3) Groups of genes that are differentially expressed in distinct groups of cells (Figure 1).

**Figure 1:**
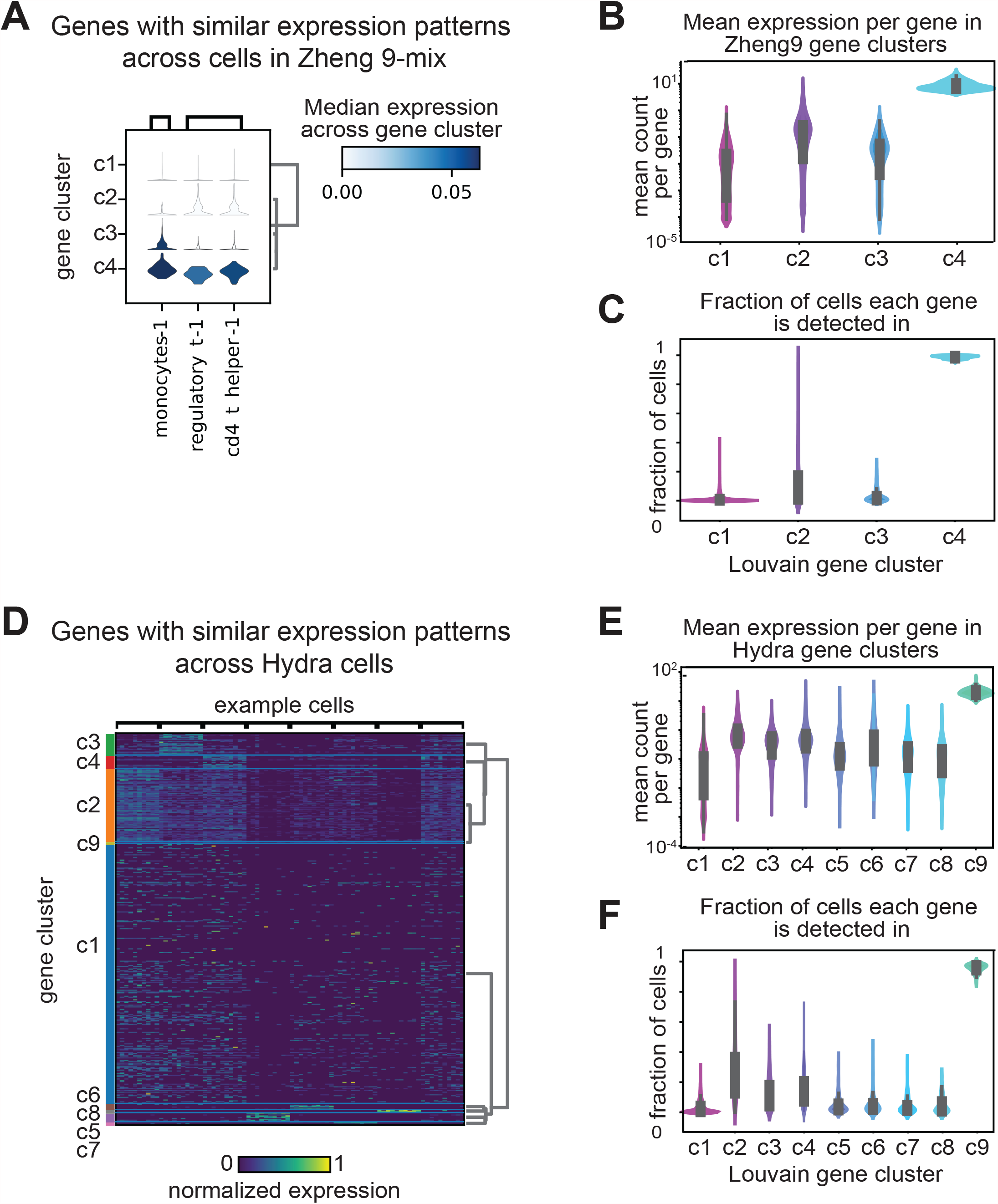
Gene clustering reveals patterns of expression variation across complex tissues and whole organisms. **A)** Violin plots depicting the estimated probability density for gene clusters in Zheng-9 lymphocytes. Each column represents a particular cell, and the violin plot is colored by the average expression level across the set of genes within the gene cluster. **B)** Kernel density estimation for the distribution of average gene expression across cells for each gene cluster group in the Zheng-9 lymphocytes. **C)** Kernel density estimation for the distribution of fraction of cells each gene in the gene cluster is identified in, for the Zheng-9 lymphocytes. **C. E)** Kernel density estimation for the distribution of average gene expression across cells for each gene cluster group in the Hydra. **F)** Kernel density estimation for the distribution of fraction of cells each gene in the gene cluster is identified in for the Hydra.

We hypothesized that genes expressed in the third, differentially expressed group are involved in the production of distinct cellular identities. In general, the physiological diversity of the sample correlated to the number of gene clusters that are only expressed in distinct groups of cells. For example, in the Zheng-9 lymphocyte data we observed one cluster of ubiquitously expressed genes (cluster 4), one cluster of sparsely expressed genes (cluster 1), and two clusters of differentially expressed genes (cluster 2 & 3) (Figure 1A,B,C). In contrast, in the Hydra data, we found six differentially expressed clusters (Figure 1D,E,F). Interestingly, in the more complex samples of mouse kidney, Hydra, and Planarian, we observed some hierarchical nature in the expression patters of differentially expressed genes, pointing to a possible subset of genes that are expressed within groups of cells that descend from a shared progenitor. For example, the Hydra gene cluster 2 is expressed across a superset of three cell-groups where each cell group expresses its own distinct subset of differentially expressed genes (Figure 1D).

Across our gene clusters, we observed a trend between the mean expression level per gene and the fraction of cells in which each gene was identified (Figure 1 B,C,E,F, S1). Genes with high mean expression levels are observed in nearly every sequenced cell. In contrast, of all the gene groups, the subset of genes which appear to be expressed at random have the lowest mean expression levels. Groups of genes that are differentially expressed have expression levels that fall between these two extremes. This observed trend between the mean expression level of a gene, and the number of cells that gene is observed within, suggests methods that relate these two metrics may be useful in identifying genes that capture real biological variation.

### A binomial model of mRNA capture identifies genes expressed in fewer cells than expected

For the analysis of scRNA-seq data, many procedures for identifying biologically varying genes have been developed. In the standard HVG feature selection approach, researchers model the mean-variance relationship (dispersion) of transformed gene count data and select those genes that are the most variable for downstream cell-type clustering. The HVG procedure depends on the assumption that variance is proportional to the mean, across the span of mean values observed in the data. Yet, in general, the mean-variance relationship depends on the distribution the sample was drawn from, and in scRNA-seq data, the distribution of particular gene expression patterns across cells is not known. Further, the variance of a gene may be underestimated when the mean counts are close to zero [Warton 2018]. To satisfy the assumptions of the HVG model and enable comparison of dispersion across genes whose means span orders of magnitude of difference, a log+1 transformation is applied to the normalized count-by-cell matrix. However, rather than achieving a variance-stabilizing effect, this transformation disproportionately increases the dispersion of genes where a greater fraction of the gene counts per cell are zeros. As a result, the HVG approach has demonstrated the capacity to enrich for a set of genes biased to have low expression values [Townes 2019]. Whether HVGs are biologically varying genes, and whether HVGs are operationally useful for cell-type clustering, has not been systematically characterized.

Other existing models assert different assumptions about the structure of gene expression and the capture process of scRNA-seq technology. The M3Drop, NBDrop, NBDisp, and Townes models all make subtly different assumptions about the relationships between the fraction of zeros per gene and the size of a cell. In the M3Drop method, the drop-out process is modeled as a kinetic process, such that there exists error around the mean counts per gene [Andrews 2019]. In the NBDrop and NBDisp method, the relationship between the fraction of zeros and mean gene count is modified by the total number of reads across each cell the gene is observed in [Andrews 2019]. These models assume that the capture efficiency changes depending on the size of a cell. Similarly, the Townes method models the mRNA capture step as a process where there is competition to be counted, essentially positing that there is a fixed number of mRNA counts per cell [Townes 2019].

In this work, we develop a different null model of variation in counts based on a binomial sampling process under simple assumptions. In our model, the observed mean expression level of a particular gene is used to develop an expectation of what fraction of cells we would find each gene in – if the gene was expressed at exactly the same level across cells (Figure 2A). This model does not require assumptions about the underlying distributions of mRNA in cells, nor the mean-variance relationship of gene expression. Based on empirical data, we propose that it is more accurate to model the binding of mRNA to UMI-coated beads as a stochastic process that is not saturated, nor affected by the size of a cell. In our model, every mRNA has an equal probability of binding to a bead, giving a binomial sampling process where each cell can be considered a trial for each species of mRNA. According to this model, the expected number of cells that should express gene *i* (*N*_*c,i*_) is given by the following expression:

**Figure 2:**
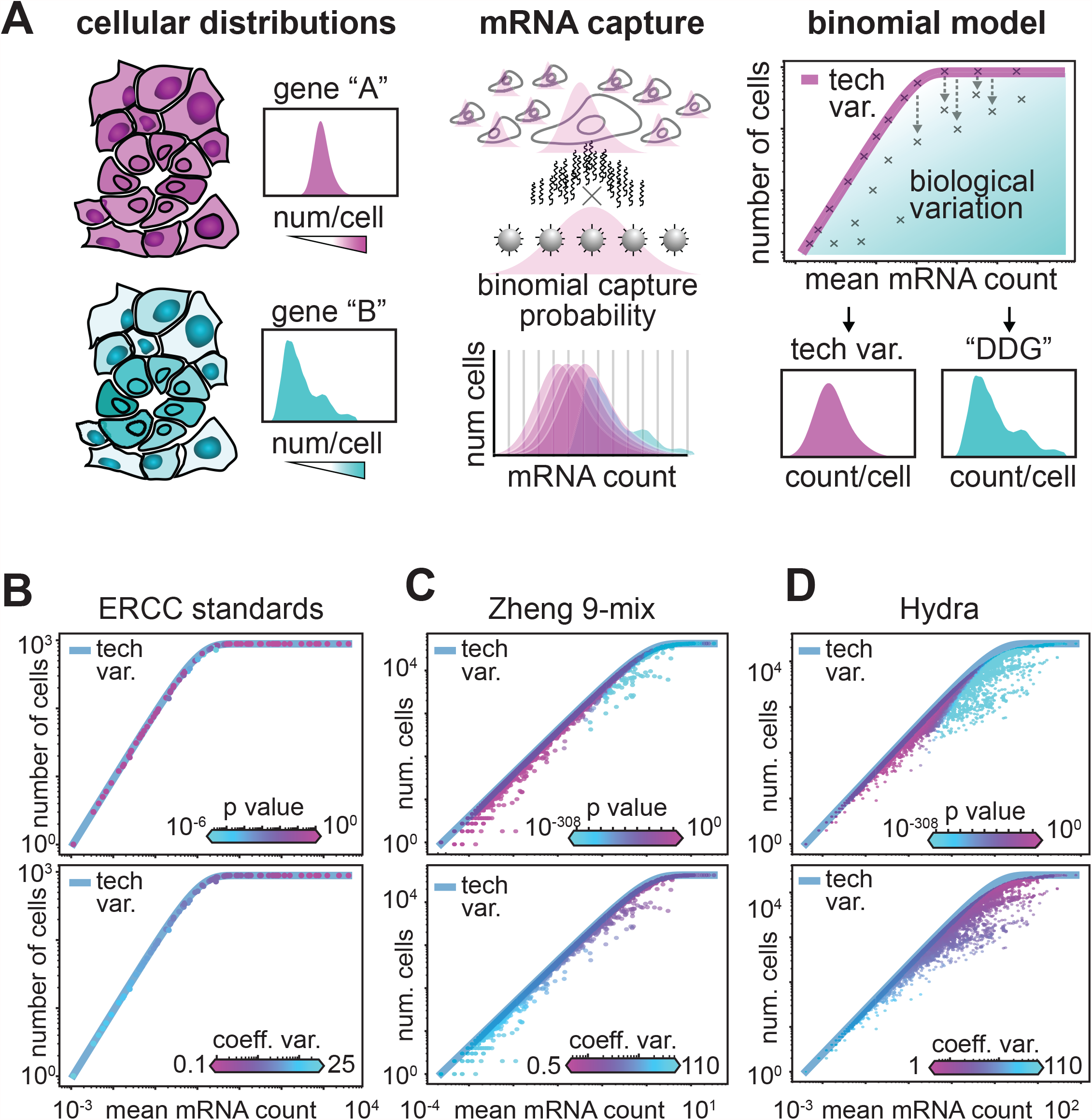
A binomial model of mRNA capture identifies genes that are expressed in fewer cells than expected. **A)** Schematic of the null model for biologically varying genes. The first panel illustrates uniformly (pink) and differentially (cyan) distributed gene expression across a sample of cells, with illustrated histograms of each genes count across the set of cells. The second panel illustrates the mRNA capture as a binomial process, where the probability of capture for each mRNA is stochastic. The third panel depicts the expected relationship of the average mRNA level and the number of cells each gene is observed in, where each ‘x’ on the graph represents a specific gene. The pink line illustrates the expected relationship if the only variation that is observed arises from the binomial sampling process. **B-D)** Scatter of average mRNA count per gene versus the number of cells each gene is identified in for three datasets: B) the synthetic ERCC data, C) the Zheng-9 lymphocytes, and D) the Hydra. In the top panel each gene is colored by the P-value computed from the DDG model, while in the bottom panel each gene is colored by the coefficient of variation.

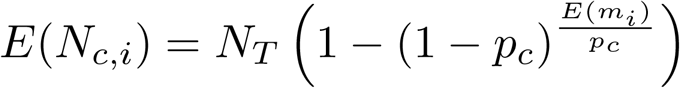

where *N*_*T*_ represents the total number of cells in the sample, *p*_*c*_ represents a fixed capture probability, and *m*_*i*_ represents the mean UMI count for each gene across all cells.

One major difference between our proposed model, and the model developed by Townes et al., is in how cell-size factors are handled. In our null model, every cell starts with an identical amount of mRNA for gene *i*, and cell-size effects can still contribute to biological variation. In other words, variation in the population that arises from cases where a cell is larger, and thus has more mRNA for gene *i* than a cell that is smaller, will deviate from the predictions of our null model. In the Townes method, the probability of capture for each mRNA is capped by its proportional expression in each cell, and natural variation in total mRNA’s per cell cannot be accounted for.

Our proposed model depends on assumptions about the source of technical variation. If the only variation observed in scRNA-seq data is variation due to sampling noise, we can expect a binomial relationship between the mean mRNA count per gene and the fraction of cells the mRNA species is observed in (Figure 2A). To test this relationship, we used scRNA-seq data created from a sample where no biological variation exists. In this experiment, scRNA-seq was performed on droplets spiked with a set of standard, synthetic ERCC control cDNA, and lacking any biological cells [Zheng 2017]. We observed that, in the absence of real biological variation, the vast majority of the spiked cDNA species fell along the expected relationship predicted by a binomial sampling process (i.e. the equation above, Figure 2B). In addition to simply predicting the relationship between mean mRNA expression and the total number of cells in which a gene is observed, we can use the model to calculate the *probability* that we would observe *N*_*c,i*_ for a given gene *i* given it’s observed mean expression level E(*m*_*i*_) (see Methods). This gives us a natural way to define a *p*-value for the null model, which is the probability of observing the same number or fewer cells expressing that gene, given *E*(*m*_*i*_). We term genes for which the observed expression pattern yields small *p*-values to be “Differentially Distributed Genes” or DDGs, since the variation in the expression pattern for that gene deviates significantly from what we would expect if the gene were expressed identically across all the cells. To correct for false identifications of DDGs due to multiple hypothesis testing, we used the Benjamini-Hochberg correction to adjust for familywise error. In the ERCC-spike in dataset, we observed that the majority of cDNA species fell on the line predicted by our simple model of the sampling process. With a familywise error rate of 1%, ∼5 out of 93 cDNA species deviate from the line of expected technical variation, suggesting that the bulk extent of technical variation in droplet-based scRNA-seq can be explained by a binomial sampling process.

We next applied our DDG model to scRNA-seq data collected from biological samples and tested the hypothesis that we would identify a greater fraction of differentially distributed genes due to underlying biological variation in gene expression. We applied our model to data collected from the Zheng-9 lymphocytes as well as the Hydra. When we plotted the mean mRNA count versus the number of cells each gene is detected in, we observe a large fraction of genes that deviate from the line predicted by our model of technical variation (Figure 2 C, D).

When each gene is colored with our *p*-value statistic, we observe that the more statistically differentially distributed genes tend to have higher mean mRNA values. Interestingly, when we color each gene by the coefficient of variation (standard deviation/mean) we observe the reverse trend: genes with low mean counts have coefficients of variation that are orders of magnitude higher than the more abundantly expressed genes (Figure 2 B,C,D). These statistics indicate that the HVG procedure may be conflating measurement noise with biological variation. Further, if the HVG and DDG methods were to be applied to raw gene counts from the same sample, the two methods may identify non-overlapping feature sets.

### Differentially distributed mRNAs exhibit different modes of variation

After identifying our sets of DDGs, we next sought to visualize the distribution of these genes across complex tissues. Because cell-identities for the Zheng-9 lymphocyte data have been annotated prior to sampling the transcriptomes, we estimated the probability density for each gene across each of the nine different cell classes. In this orthogonally annotated data, we observed three general patterns of gene expression that can occur independently from mean expression value. 1) For genes with non-significant *p*-values, we find uniform distributions both across and within different cell types (Figure 3B). This class of genes are those genes whose expression levels do not vary more than what would be expected from a binomial capture process. These genes also tend to include, but are not limited to, genes that have mean expression values that are close to zero (Figure 3A, Figure 4). 2) In the Zheng-9 lymphocytes, we observe DDGs that are differentially expressed in specific cell types (Figure 3C). For example, out of the 9 annotated cell types, the gene GNLY is only highly expressed in Natural Killer cells, as it encodes a specialized protein with antimicrobial activity, called granulysin. 3) We also find DDGs that quantitatively vary both across and within cell types (Figure 3D). The proteins encoded by these genes include general cell-function proteins such as ribosomal proteins and S100 family calcium binding proteins, as well as proteins with established immune function. For example, we observed increasing counts of the GZMM serine protease across activated lymphocytes. We also observed different levels of the CD52 peptide that is associated with the mobility of lymphocytes, as well as other cytokine receptor and antigen associated genes.

**Figure 3:**
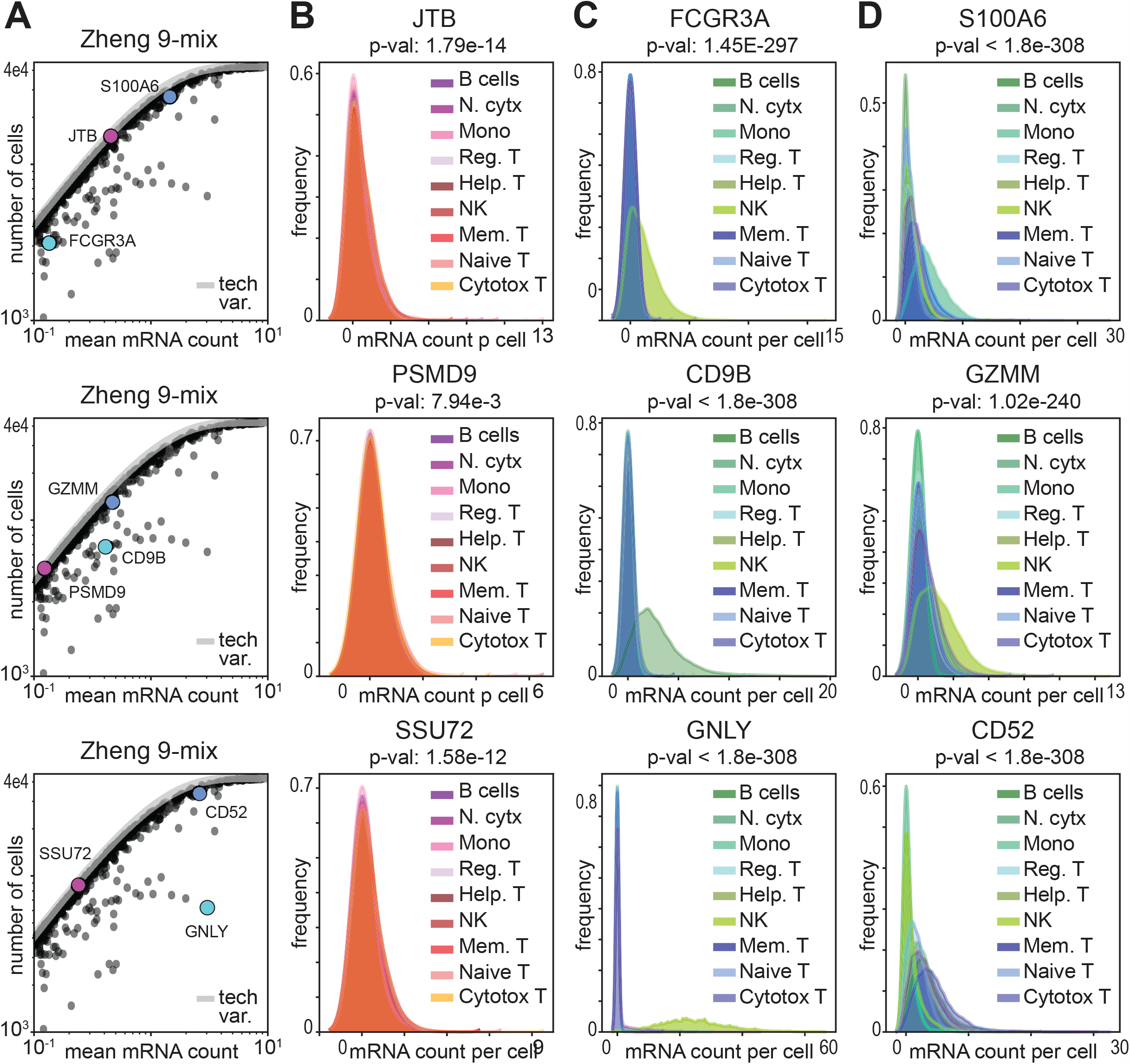
Identification of differentially distributed mRNAs reveals different modes of expression variation. **A)** Scatter of average mRNA count per gene versus the number of cells each gene is identified in for the lymphocyte data, cropped to highlight the three example genes depicted across the row. **B-D)** Kernel density estimation for the distribution of gene counts across cells in the lymphocyte data. The distribution of each gene is plotted for each cell type separately. In each column three example gene distributions are depicted to show B) genes that are not significantly differentially distributed, C) genes that are differentially expressed in specific cell types, and D) genes that quantitatively vary both across and within cell types.

**Figure 4:**
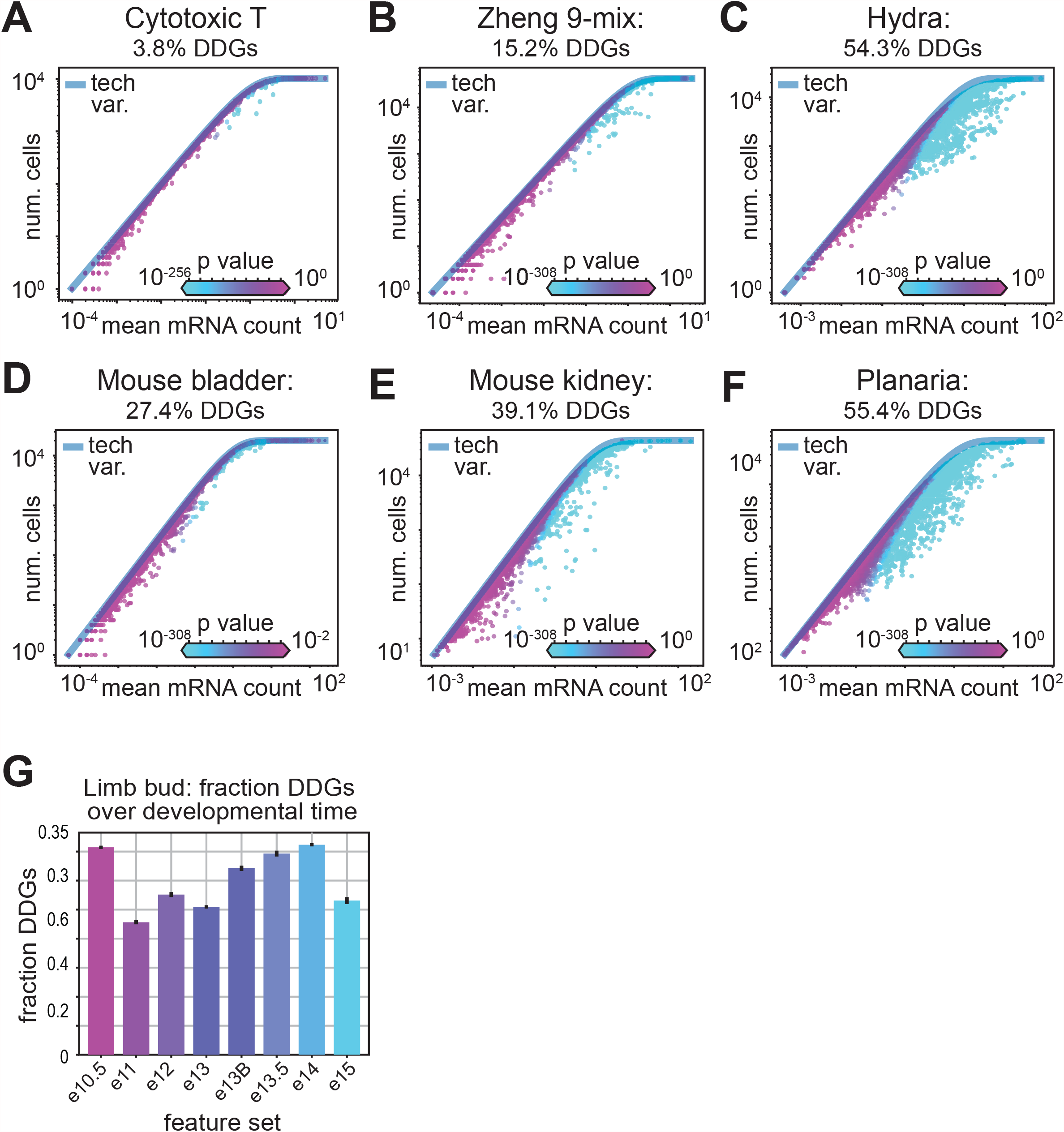
Greater tissue complexity produces a greater fraction of differentially distributed genes. **A-F)** Scatter of average mRNA count per gene versus the number of cells each gene is identified in for the lymphocyte data, where each gene is colored by the computed p-value. The fraction of significantly distributed genes is indicated for A) cytotoxic T cells, B) the Zheng-9 lymphocyte mix, C) hydra, D) mouse bladder, E) mouse kidney, and F) planaria. **G)** Mean fraction of DDGs over developmental time in mouse limb bud data. Error bars show confidence interval around the mean, estimated using the 10 re-sampled sets.

### A natural method for determining the number of genes in an scRNA-seq feature set

Several feature selection procedures use an arbitrary cutoff for determining the number of biologically informative genes. In the HVG method, typically 2,000-5,000 of the most variable genes are chosen to create the cell-by-gene distance matrix upon which cell-type clustering is performed. Arbitrary cutoffs are also imposed by the Townes and NBDisp method, while the M3Drop and NBDrop models use a false discovery rate-adjusted test statistic. Rather than choosing an arbitrary number of genes for an analysis, we hypothesized that we should expect the number of biologically informative genes to increase with the number of different cell identities within a sample. To test this hypothesis, we calculated the fraction of DDGs over samples with increasing tissue complexities.

For tissues with more distributed physiologies, we recovered a greater fraction of differentially distributed genes. For example, we found a fraction of only 3.8% DDGs in an isolated sample of cytotoxic T cells, compared to 15.2% DDGs across the Zheng-9 mix (Figure 4A,B). Similarly, in the mouse bladder, where three major cell types are expected, 27.4% of the genes are differentially distributed, while we find 39.1% DDGs in the more physiologically diverse mouse kidneys (Figure 4D,E). When we calculated the fraction of DDGs across scRNA-seq data collected from whole multicellular organisms, we observed 54.3% DDGs in the hydra and 55.4% DDGs in Planaria (Figure 4C,F). We then repeated this experiment using the two other feature selection models where a natural test statistic determines the number of significant genes for each sample. Interestingly, in the NBdrop method, we observed a similar trend as in the DDGs, where the number of genes increased as a function of tissue complexity (Figure S4B). However, the overall number of genes was less for each sample, and the kidney sample deviated from the trend with higher numbers of significant genes than the two whole-organism samples. In contrast, the M3drop method produced a fraction of significant genes that decreased with increasing tissue complexity, contradicting our knowledge of gene expression variation across complex tissues (Figure S4C).

We next tested the hypothesis that the fraction of differentially distributed genes increases over developmental time. For this experiment, we used scRNA-seq data collected from mouse limb buds spanning 10.5-15 weeks in embryonic development [He 2020]. To ensure equivalent statistical power, we randomly sampled 5,000 cells from each time point ten times, calculated the fraction of DDGs, and generated confidence intervals across our samples. We found that as developmental time progressed, the fraction of DDGs generally increased, corroborating our hypothesis that diversity of cell types corresponds to diversity in gene expression (Figure 4G). However, we found that the earliest and latest limb bud sample (e10.5 and e15) deviate from the expected trend. To account for this deviation, we plotted quality control metrics across each of the samples (Figure S4 D,E,F). We found that the distribution of total UMI counts across cells within the e10.5 and e15 sample also deviates from the observed trend across the samples, and positively correlate to the fraction of DDGs recovered (Figure S4 F).

The fraction of DDGs recovered also depends on how many cells are used to generate the observations. To evaluate how the power of our model depends on the number of cells observed, we took an empirical approach to account for the diversity of cell types that are observed in real biological data. Because we have FACs labels for lymphocyte data, we can approximate a set of “real” DDGs using a supervised approach. Specifically, we calculated the set of differentially expressed genes across these nine groups, using the Wilcoxon rank-sum test. We used the same FDR cutoff for the Wilcoxon rank-sum genes as we used for defining significant DDGs. We randomly sampled each of the nine types of lymphocytes over a range of 100 to 5,000 cells, calculated the set of DDGs, and computed the overlap between the “real” DDGs obtained from the Wilcoxon rank-sum test and the DDGs predicted by our model, as a function of increasing cell number. First, we find that the number of DDGs recovered increases linearly as a function of number of cells (Figure S4 G). When we calculate the overlap between the predicted DDGs and the true DDGs, we find that as we increase the number of cells, the number of true DDGs recovered begins to saturate (Figure S4 J). Together, these results suggest that the power of the DDG model depends linearly on the number of cells observed, yet the number of biologically variable genes is finite, and with an increasing number of cells, the DDG model can recover an increasing fraction of true variable genes.

### Preservation of variance structure during feature selection

After validating our DDG model using technical and biological controls, we proceeded to comprehensively characterize different feature selection methods. In the standard scRNA-seq analysis workflow, feature selection is motivated by the idea that dimensionality reduction can remove axes of variation that arise due to sampling noise, and thus improve the identification of cells that vary in similar and biologically informative ways. Feature selection typically implies a 10-20-fold reduction of genes that are used to represent cells, before the dimensionality is further reduced by principal component analysis, and clustering algorithms are applied. However, the extent to which various feature-selection models can preserve the variance structure of high-dimensional data, or of real biological variation, has yet to be established.

To evaluate whether cell neighborhood topologies can be preserved by various feature selection methods, we calculated the distortion induced to cell neighborhoods by dimensionality reduction using a metric called the Average Jaccard Distance (AJD). The AJD is defined as the per-cell average of the difference between the overlap and total set of k-nearest cell-neighbors in the high-dimensional space, compared to the k-nearest neighbors in the reduced dimensional space (Figure 5A) [Cooley 2020]. If the AJD = 0, all cell-neighbors are the same, and no distortion is induced by the dimensionality reduction. In contrast, if the AJD = 1, dimensionality reduction has permuted all of the neighbors, such that none of the original k-nearest neighbors in the high-dimensional data remain in the reduced-dimensional projection.

**Figure 5:**
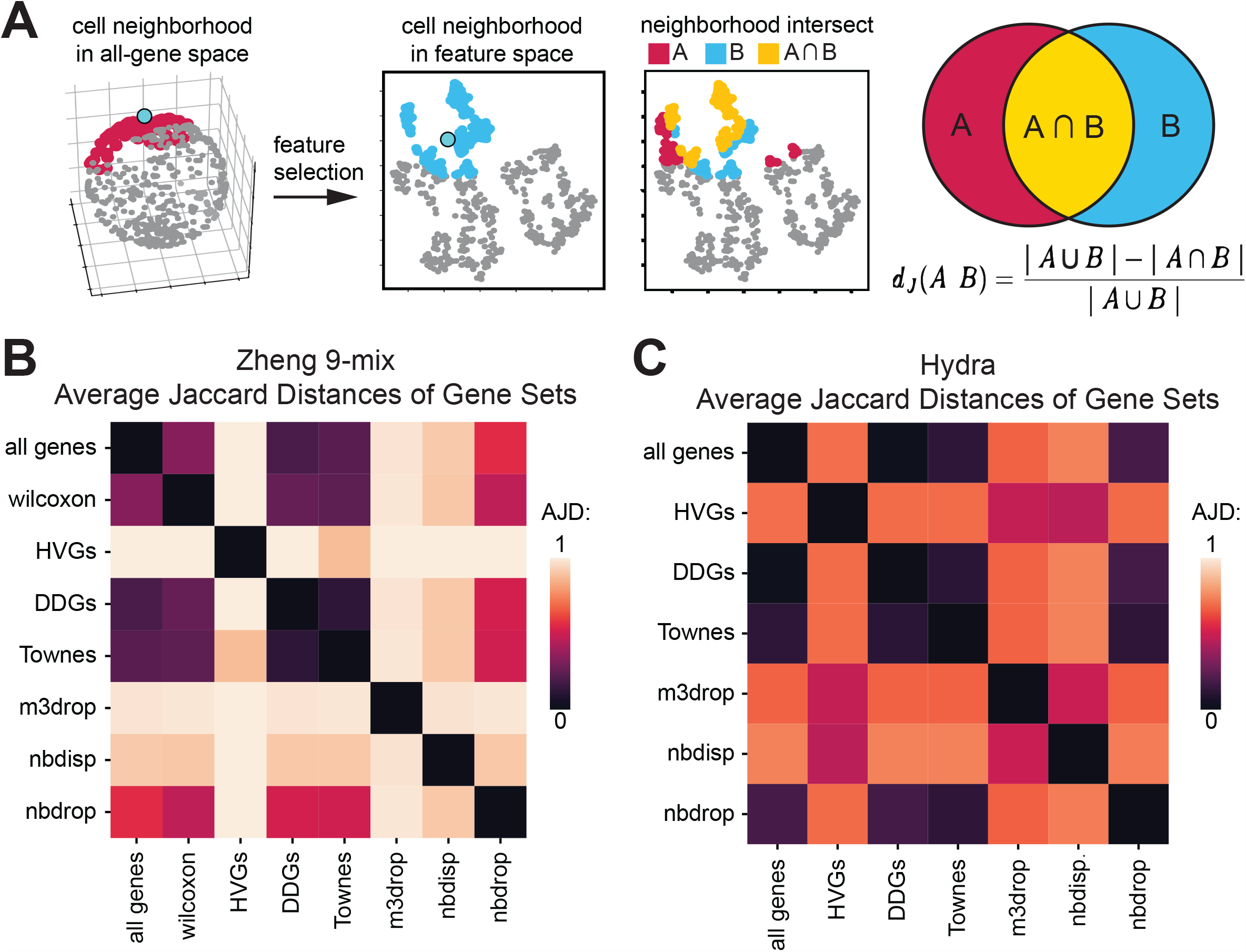
DDGs preserve the structure of variation in high-dimensional scRNA-seq data. **A)** Schematic of the Average Jaccard Distance as a measure of distortion induced by dimensionality reduction **B**,**C)** Heatmap of pairwise Average Jaccard Distances for each feature set for the B) Zheng-9 lymphocyte mix and C) Hydra

Using the data where all genes are included as a reference, we calculated the neighborhood distortion induced when various feature selection methods were applied to the Zheng lymphocytes (Figure 5B). Using our supervised set of differentially expressed genes, determined by applying the Wilcoxon rank sum test across the FACS labeled cell types, we first established a minimum expectation of distortion induced by feature selection. We found that when cell-neighborhoods were calculated using 20 nearest neighbors and the supervised set of 2,061 genes, the variance structure of the full 20,000 gene set was largely preserved. The Wilcoxon set produced an AJD of only 0.34. In comparison, when the HVGs were used as the basis for the dimensionality-reduced space, the AJD was 1.0, suggesting that all of the biological variation expressed in the local neighborhood structure has been lost. In contrast, when we calculated the distortion induced by the DDG set, we found a AJD of only 0.19, indicating the DDG set preserves the high-dimensional neighborhood structure better than the Wilcoxon genes. This slight improvement of the DDGs over the Wilcoxon genes may arise from the inability of the Wilcoxon test to identify genes that quantitatively vary within cell types or across more than one cell type. We next compared the preservation of variance by DDGs to that of other models. We found that the other two binomial-based models comparable to the preservation of variance by DDGs, yet the NBdrop and Townes method induces slightly greater distortion with values of 0.56 and 0.23, respectively. In contrast the M3drop procedure more significantly permutes the neighborhood structure relative to both the all-gene and Wilcoxon-gene distance matrices, giving an AJD of 0. 98. These trends were also observed across Hydra – relative to the high-dimensional data, HVGs greatly permutated the neighborhood structures, while DDGs induce almost no distortion (Figure 5C).

### Recovery of differentially expressed genes by different feature selection methods

We next evaluated whether various feature selection models could recover the specific set of differentially expressed genes that were generated using the supervised Wilcoxon rank-sum test on the FACS-labeled lymphocytes. Out of all the feature selection methods tested, the DDG set recovered the largest fraction of the Wilcoxon genes, specifically sharing 1343 out of a total of 2061. In contrast, HVGs only recover 689 (Figure 6A). We calculated the Jaccard Index to account for the proportional overlap as different feature selection methods give feature sets of different sizes when applied to the same sample data. Again, DDGs had the highest proportional overlap with the Wilcoxon genes, while the HVGs had the smallest overlap (Figure 6B).

**Figure 6:**
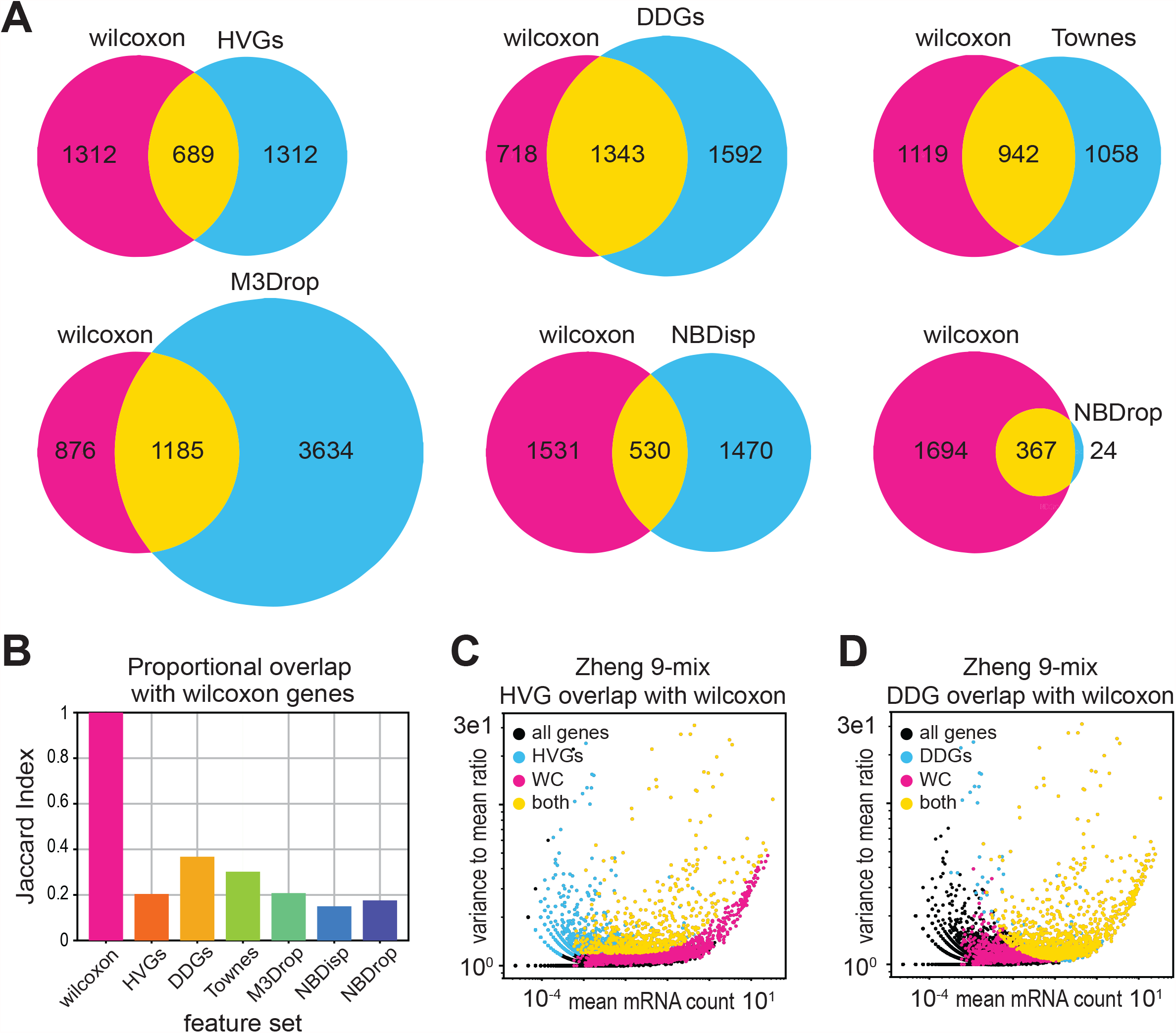
DDGs recover supervised gene-expression differences of FACS-labeled lymphocytes. **A)** Venn diagrams of set overlap for each feature set and the set of genes identified in the Zheng-9 lymphocyte mix using the Wilcoxon rank sum test. **B)** Jaccard index for each feature set illustrating the size-adjusted overlap with the Wilcoxon gene set in the lymphocyte mix. **C**,**D)** Scatter of the mean mRNA count versus dispersion per gene, colored by set membership for feature sets in the lymphocyte data. C) compares HVGs and Wilcoxon genes while D) compares DDGs and Wilcoxon genes.

Next, to develop intuition about what kind of bias may be introduced by various feature selection models, we plotted the mean mRNA count against the coefficient of variation or “dispersion” (the variance to mean ratio), for each gene in the Zheng lymphocyte data. Here, each gene is colored by its membership to the supervised Wilcoxon set, a particular feature selection set, or membership to both the Wilcoxon and the feature set (Figure 6C,D). When we compared the Wilcoxon genes to the HVGs, we observe that HVGs tend to have lower mean expression, but high dispersion (Figure 6C). This finding corroborates the previous hypothesis that the HVG procedure produces a biased output of the most variable genes towards a set of genes with low mean expression but high variance. In contrast, DDGs fail to identify Wilcoxon genes with lower expression levels, potentially due to a lack of power at lower mean expression levels.

We then compared the DDG model to the other more similar, binomial-based methods (Figure S6 B,C,D). Interestingly, the genes included in the DDG set are similar to those in Townes, yet each feature set has a unique subset of genes. Additionally, the Townes method tends to select genes that have higher expression means than the DDGs, suggesting a lesser ability to identify differentially expressed genes with lower expression values (Figure S6C). Meanwhile, the NBdrop method produces a set of genes that are a subset of the DDGs, yet have much fewer genes in comparison (Figure S6D). The NBdrop method also tends to select the subset of DDGs that have high dispersion values, again indicating a weaker power in the ability to resolve differentially expressed genes. Taken with the previous findings, these results suggest that compared to other methods, our binomial model with simpler assumptions can identify a greater fraction of true biologically varying genes.

### Recovery of FACs labels by different feature sets

We next sought to test the operational goal of feature selection which is to enable recovery of physiologically similar cells that have similar transcriptional profiles, despite the sparseness of scRNA-seq data that results from low capture probabilities. To evaluate which feature set is most informative for this task, we used the FACS-labeled lymphocytes, where biological identities are determined *a priori* based on historically-established notions of major immune cell types. In this experiment, we apply the standard approach of Louvain clustering to the dimensionality-reduced data and evaluate how well each feature set can recover the original FACS labels.

For each feature-selected group, we titrated the Louvain resolution parameter from 0.1 to 1, calculated the adjusted rand index (ARI) to compare set membership between Louvain clusters and FACS labels across all cells, and plot the best score per group [Rand 1971]. In the first experiment, we found that when using the raw UMI counts as the basis for the cell-cell distance calculations, none of the feature selection groups were able to achieve an ARI score greater than 0.5, indicating that cells were not being partitioned into their historically established cell-type groups (Figure S7G). Yet, rather than using raw UMI counts as the basis for cell-type clustering, the standard approach to scRNA-seq analysis performs cell clustering on feature sets where gene sets are first CPM and log+1 transformed, and then the cell-by-gene data matrix is further reduced by principal component analysis (PCA). The application of PCA to the feature selected data is motivated by the idea that reducing the sparseness of the data while retaining the variance structure will enhance the performance of the Louvain clustering algorithm. To test this idea directly, we performed Louvain clustering various steps in the scRNA-seq analysis pipeline by using raw counts, PCA-transformed raw counts, and Log+1-CPM-PCA transformed counts. We found that using PCA-transformed counts only marginally improved the performance of various feature selection sets, whereas the full log+1-CPM-PCA transformation increased the ARI scores of the all genes, or binomial feature sets to slightly greater than 0.6 (Figure S7H,I).

We then visually inspected the tSNE projections where cells were colored by either their original FACS labels or their Louvain identified clusters, to identify where the Louvain algorithm was failing to appropriately partition cell clusters [van der Maaten 2008]. We observed that none of the assayed feature sets were able to recover the appropriate FACS labels for the subsets of related T-cell lineages (Figure S7A-F). This observation corroborates the existing idea that there might not exist a transcriptional basis that can separate these related T cell lineages into distinct groups that map to distinct physiological roles [Omilusik 2017, Henning 2018, Kiner 2021]. Based on these observations, we next performed a second FACS-label recovery experiment, where we merged the FACS labels from two subsets of specialized T-cell lineages into two distinct supersets. Specifically, we combined naïve cytotoxic cells, cytotoxic T cells, and memory T cells into one set, and naïve T, helper T and regulatory T cells into another set (Figure S7A-I). We then proceeded to evaluate the performance of different feature sets using a more granular approach where ARI scores were not dominated by differences in the capacity of feature sets to recover differences within these distinct T cell lineages.

Several gene sets were able to recover a large majority of the initial FACS labels, giving ARI scores over 0. 9 (Figure 7). These gene sets include all genes, the Wilcoxon genes, the DDGs and the Townes genes. We found that log+1-CPM-PCA transformed counts again perform slightly better than no transformations or only PCA transformation, potentially by permutating neighborhood structure in a way that is more compatible with the Louvain algorithm (Figure 7G,H,F). Interestingly, the variance-based approaches, including the HVG set, failed to produce an ARI of greater than 0.6 (Figure 7G,H,F), even with the minor T-cell lineages merged into major groupings. We then inspected the tSNE projections for the HVG set, where cells are labeled according to either their FACS labels (Figure 7B) or their Louvain groups (Figure 7E). The comparison of these plots suggests that the separatrix that distinguishes the two separate T cell lineages in the all genes and DDG set (Figure 7A,C) appears to be lost in the HVG set. Thus, when Louvain algorithm is applied to the HVG set, it fails to recover a large proportion of the original FACS labels.

**Figure 7:**
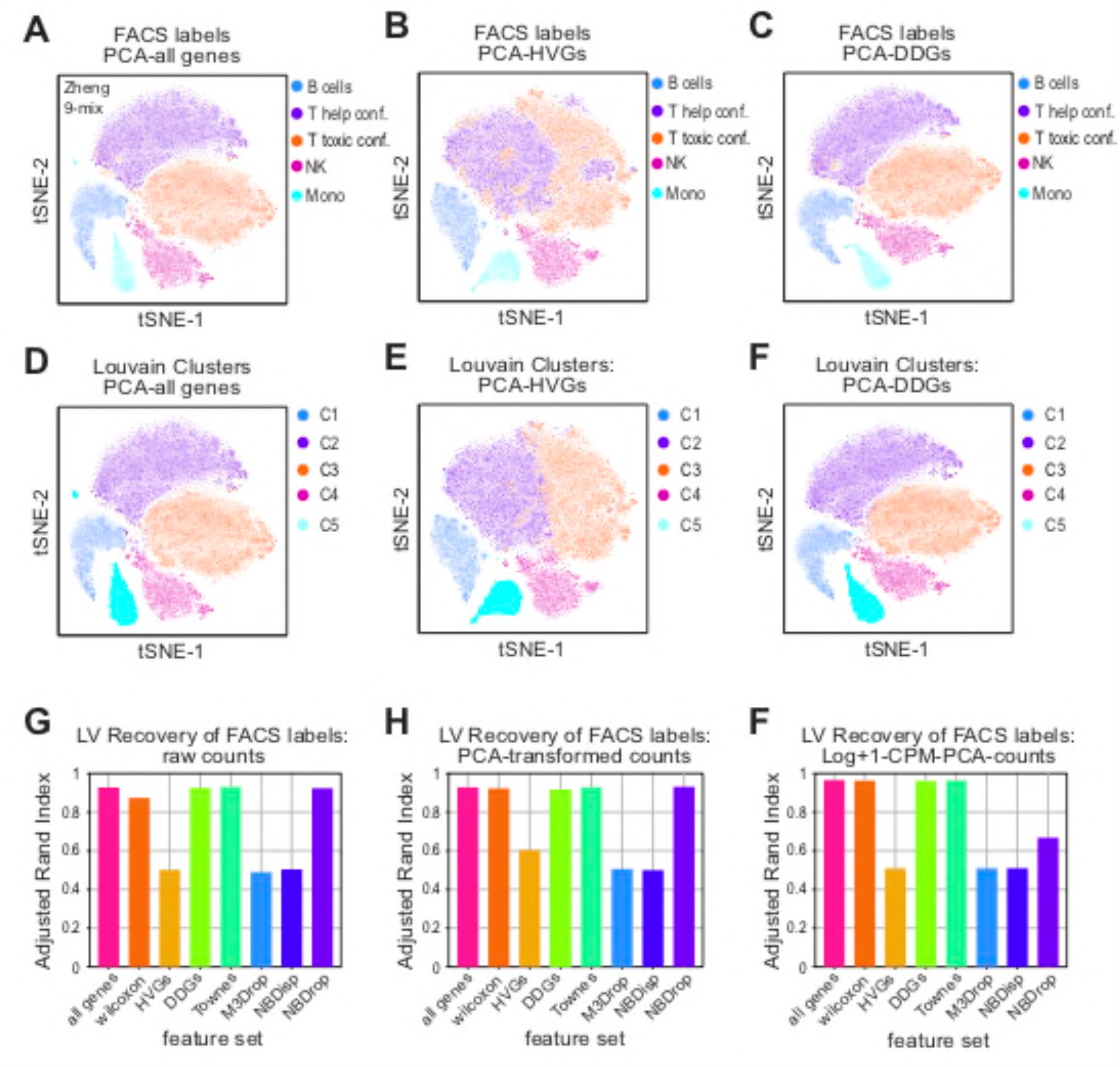
DDGs recover cellular identities of FACS-labeled lymphocytes. **A-F)** tSNE projections for dimensionality-reduced data from the Zheng-9 lymphocyte mix. Principal component analysis was used to reduce (A,D) all genes, (B,E) HVGs, or (C,F) DDGs. Cells are either colored by original FACs-based set membership, where T-celI lineages were merged into two super sets (A-C), or Louvain cluster membership (D-F). **G-F)** Quantification of FACs label recovery by various methods of dimensionality reduction across different feature sets. Louvain clustering was performed across a titration of resolution parameters, cluster labels were compared to original FACs labels where T-cell lineages were merged into two super sets, and the highest adjusted rand index is plotted when G) raw UMl counts, H) PCA-transformed counts, or F) Log+1-CPM-PCA-counts were used as a basis.

## Discussion

A major challenge in the study of single cell biology is understanding how variation in gene expression maps to cell physiology. Single-cell RNA sequencing seeks to overcome this challenge by enhancing the resolution at which transcriptional variation can be measured, yet the interpretation of these datasets is subject to the precise mapping challenge it seeks to overcome. The dominant approach to interpreting transcriptional variation relies on clustering groups of transcriptionally similar cells, and then comparing differences in gene expression patterns between the groups. Ideally, unsupervised clustering can partition groups of cells that vary in similar ways into biologically meaningful groups. Yet defining the most appropriate notion of transcriptional similarity and the best way to delineate these cells into cell-type classes remains an ongoing challenge [Qi 2020].

Nonetheless, an operational goal of scRNA-seq studies is to identify genes that are related to biological variation. However, the nature of scRNA-seq data poses challenges for this goal, as the low probability and stochasticity of the scRNA-seq capture process generates sparse, noisy, and extremely high-dimensional gene count data. These raw UMI measurements can conflate biological and technical variation and may create challenges for existing unsupervised clustering algorithms. Therefore, a critical step in the analysis of this data is developing models of measurement noise that allow researchers to accurately distinguish which axes of variation are biological and which arise due to technical noise in the measurement.

Within the domain of scRNA-seq analysis, evaluating which null model of biological variation requires empirical support, where biologically meaningful transcriptional variation can be compared to the output of feature selection models. In this work we propose a feature selection model with minimal assumptions. While our model is similar to the NBDrop and Townes methods, our model relaxes constraints about how cell size affects the mRNA capture process, and treats every mRNA as having an equal probability of being captured. We benchmark how our model performs compared to other popular methods, by using data where known biological variation has been pre-established. Specifically, the Zheng lymphocyte data, where cells had been sorted into major cell type classes and annotated with FACs labels prior to sequencing, enables us to directly map biological variation to transcriptional variation. In addition, because the Zheng lymphocyte data has a relatively large number of cells in the data set, we were able to test model performance using a several rigorous metrics.

We demonstrate that, compared to other approaches, our DDG method produces the most accurate mapping of established physiological variation to dimensionality-reduced transcriptional variation. First, the DDG method assigns a *p*-value during the feature selection process, and provides a natural cutoff by which the number of features can be determined after correction for multiple hypothesis testing. We show that this approach is the only method tested that allows us to reliably recover an increasing number of biologically varying genes when applied to samples with increasing tissue complexity. Second, DDGs preserve high-dimensional variation in the neighborhood structure of raw count-by-cell data, suggesting the major axes of transcriptional variation are retained in the reduced dataset. Further, using the orthogonally-annotated lymphocyte data, we find that the DDG feature selection method best recovers a set of supervised differentially expressed genes, and the DDG-set provides a basis for unsupervised clustering that enables recovery of the original FACS cell-identity labels.

Taken together, these findings support the use of DDGs in improving the standard scRNA-seq analysis pipeline. Our finding that HVGs severely distort the neighborhood structure of the high-dimensional data and perform poorly across all other tested metrics, corroborate the existing hypothesis that HVGs create an almost arbitrary basis for cell type clustering. We argue that the HVG method biases feature selection toward those genes with low expression values, and that this popular method is not appropriate for dimensionality reduction or feature selection for scRNA-seq data. Moreover, our cell-clustering results suggest feature selection only offers a marginal improvement over clustering cells using the full, raw count by cell matrix. Nonetheless, dimensionality reduction using DDG feature selection can offer an advantage when computational power is limited. DDGs also offer a viable dimensionality reduction approach for other computationally intensive techniques such as manifold learning, when PCA may not be the most appropriate method to identifying manifolds of transcriptional variation at the single cell level.

DDGs identify genes that are differentially distributed in their expression both within a specific subset of specialized cell types, as well as those that vary more continuously both within and across transcriptionally similar groups of cells. Thus, the subset of DDGs provides greater information than standard differential expression tests, which are not designed to identify genes that quantitatively vary across multiple cell classes, or within a particular group. As such, the standard scRNA-seq analysis pipeline overlooks potentially interesting manifolds of transcriptional variation. Overall, this selective vision highlights a more general struggle in biological research. Classifying cells into discrete types can be operationally useful for studying changes in gene expression. However, to what extent transcriptional identity maps to discrete objects, or whether physiologically distinct cell-types correspond to well-separable regions in transcriptional space, remains underexplored. The artificial boundaries that define abundance and composition of different cell types, paired with the oppositional logic that highlights the presence or absence of specific genes, limits our perspectives on cell physiology. Future work might utilize the DDG method to retain informative axis of biological variation, and enhance our comprehension of cell variation by permitting a more dynamic framework by which we understand cell identity transitions.

## Methods

### Binomial model of the mRNA capture process

The basis for our null model of variation due to technical noise is quite simple. When run through most scRNA-seq pipelines, cells are lysed in the presence of beads that capture mRNA molecules and then label those mRNAs with both cell-specific barcodes and Unique Molecular Identifiers during the subsequent PCR steps. In our model, we imagine that every mRNA molecule in the cell has the same probability of being captured during this process; we refer to this probability as *p*_*c*_. Say that, for a given gene *i* in cell *j*, there are a total number of *s*_*i,j*_ copies of mRNA of that gene in that cell (we use the variable *s* here to denote the “starting” number of molecules). Under this model, the observed number of mRNAs we obtain in the data will follow a simple binomial distribution:

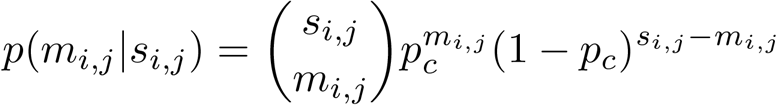

where *m*_*i,j*_ is just the observed number of UMI counts for that mRNA in cell *j*.

### Expected number of cells expressing a gene i

The first step in developing our approach involves determining what the binomial sampling model described would predict under the very simple null model that there is actually no biological variation in gene *i* within the original sample. In other words, imagine that we had a situation where *every single cell* started out with the same number of copies of mRNA for that gene (i.e. *s*_*i,j*_ *= s*_*i,k*_ ∀ *j,k*). Of course, even in this scenario, there would be some variation in the resulting scRNA-seq data, since the sampling process is binomial and there would be some differences between cells simply because the capture process is stochastic. Our ultimate goal is to identify genes whose expression pattern is inconsistent with this null expectation; in other words, genes whose variation within the data *cannot be explained* purely on the basis of the stochastic process of mRNA capture during the experiment.

In order to proceed, we first consider what this model would predict in terms of the relationship between the *number of cells* that express a gene *i* (i.e. the number of cells with *m*_*i,j*_ > 0, call this *N*_*c,i*_) and the average expression of that gene in the population *E*(*m*_*i*_) (note here that we drop the index *j* since we have averaged over all the cells). The reason for doing this is the simple fact that the binomial model above immediately suggests a simple relationship between these two observations. Call the total number of cells within the dataset *N*_*T*_. The first step is to determine the value of *s*_*i,j*_ that we would expect given the observed data. It is clear that the maximum likelihood estimator of this parameter is just *E*(*m*_*i*_)/*p*_*c*_, or just the observed average number of mRNAs of that gene across the cells in the population divided the capture probability.

Now, we imagine that each of our *N*_*T*_ cells is subjected to the binomial sampling process independently, all starting from *E*(*m*_*i*_) mRNA molecules. The average number of cells expressing this gene should be:

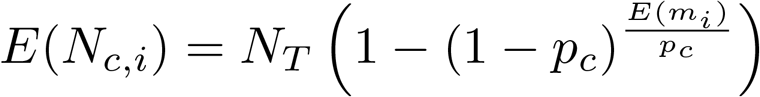

note that the term within the parentheses is just the probability that you will capture 0 mRNAs from the cell, given *s*_*i,j*_ = *E*(*m*_*i*_); in other words, this is just *p*(0|*E*(*m*_*i*_)). We multiply this probability by the total number of cells to obtain the expectation.

#### p-value calculation under the null model

We calculate a *p*-value under this null model for a given gene *i* using a similar approach. Essentially, if we know the probability that we should observe any given cell with a count of 0 under the null model, then we can view the whole scRNA-sequencing data set as a different kind of binomial trial. Each cell is subjected to sequencing, and it either expresses the gene (*m*_*i,j*_ > 0) or it does not (*m*_*i,j*_ = 0). For simplicity, define:

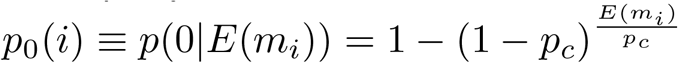

as the probability of having a cell with 0 copies of gene *i*. From this definition, it is easy to see that the probability of observing *N*_*c,i*_ (the number of cells expressing gene *i*) is just:

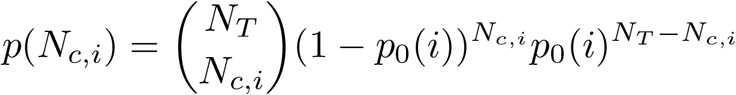

Obtaining the *p*-value for gene *i* is then a simple matter of summing this probability for all values of *N*_*c,i*_ less than or equal to the observed value in the data.

