## Supplementary Figures for "Binomial models uncover biological variation during feature selection of droplet-based single-cell RNA sequencing"

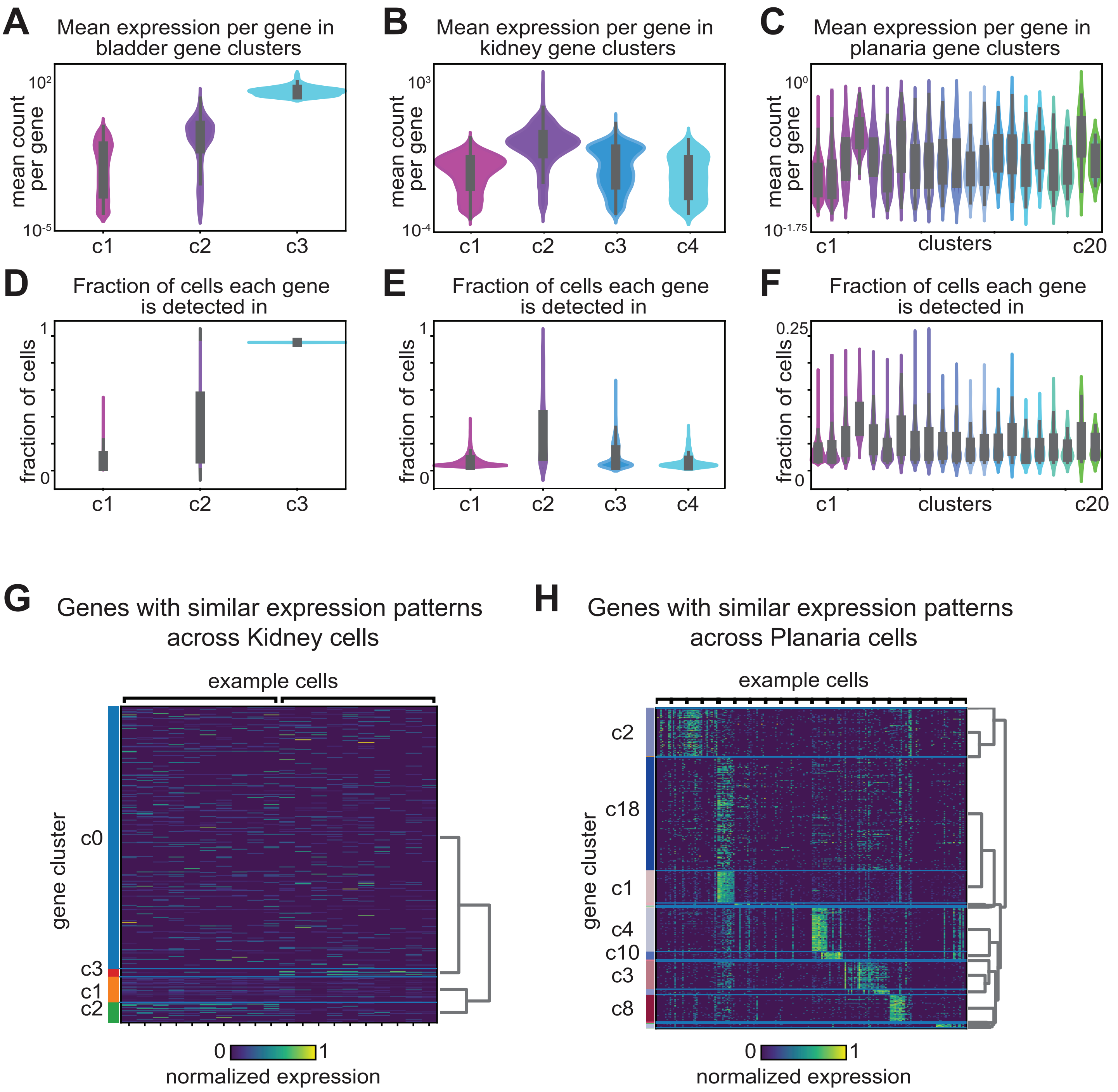


**Figure S1: A-C)** Kernel density estimation for the distribution of average gene expression across cells for each gene cluster group in A) mouse bladder, B) mouse kidney, and C) Planaria. **D-F)** Kernel density estimation for the distribution of fraction of cells each gene in the gene cluster is identified for D) mouse bladder, E) mouse kidney, and F) Planaria. **G-H)** Heat map depicting the normalized expression level of genes grouped by gene cluster membership in the G) kidney and H) Planaria. Each row represents a gene and column represents a particular cell, and the entry is colored by the normalized expression level

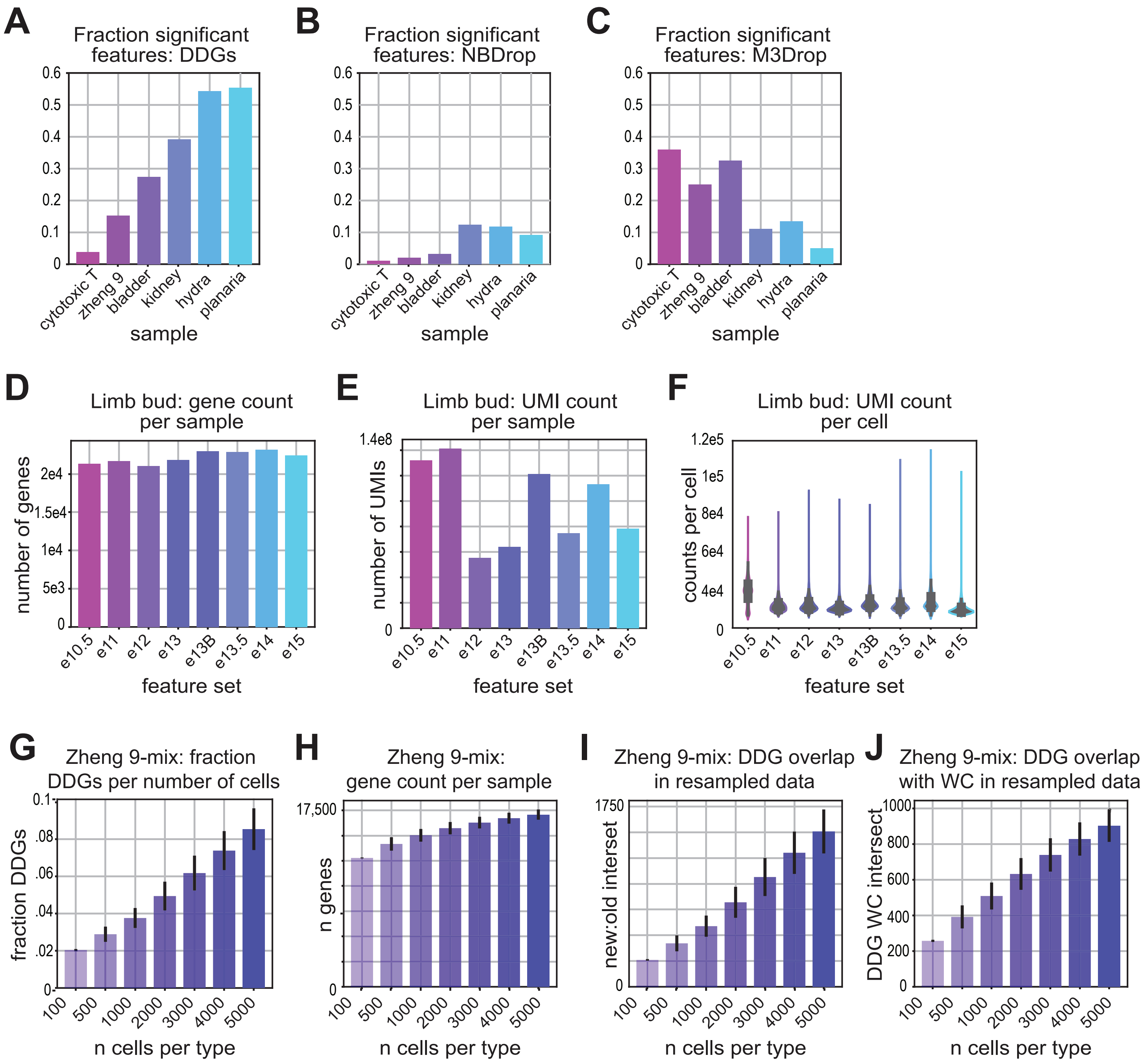


**Figure S4: A-C)** Mean fraction of significant features over increasing tissue complexity using A) the DDG method, B) the NBDrop method, and C) the M3Drop method. **D-F)** Quality control metrics for the limb bud data. A) Mean total number of genes for each sample, with error bars depicting confidence intervals around the mean. B) Mean total number of UMI counts for each sample, with error bars depicting confidence intervals around the mean. C) Distribution of UMI counts per cell in each limb bud sample. **G-J)** Empirical power estimations for the DDG method using the Zheng-9 lymphocyte mix, where a new set of DDGs was computed with increasing sample size. G) Mean number of DDGs as a function of increasing number of cells per cell type in the Zheng-9 lymphocyte mix. H) Mean total number of genes as a function of increasing cell number per sample. I) Mean overlap of new DDG sets with original DDG set calculated from the full, 5k cells per type Zheng-9 lymphocyte data. J) Mean overlap of new DDG sets with original Wilcoxon set of differentially expressed genes, as a function of number of cells per cell type.

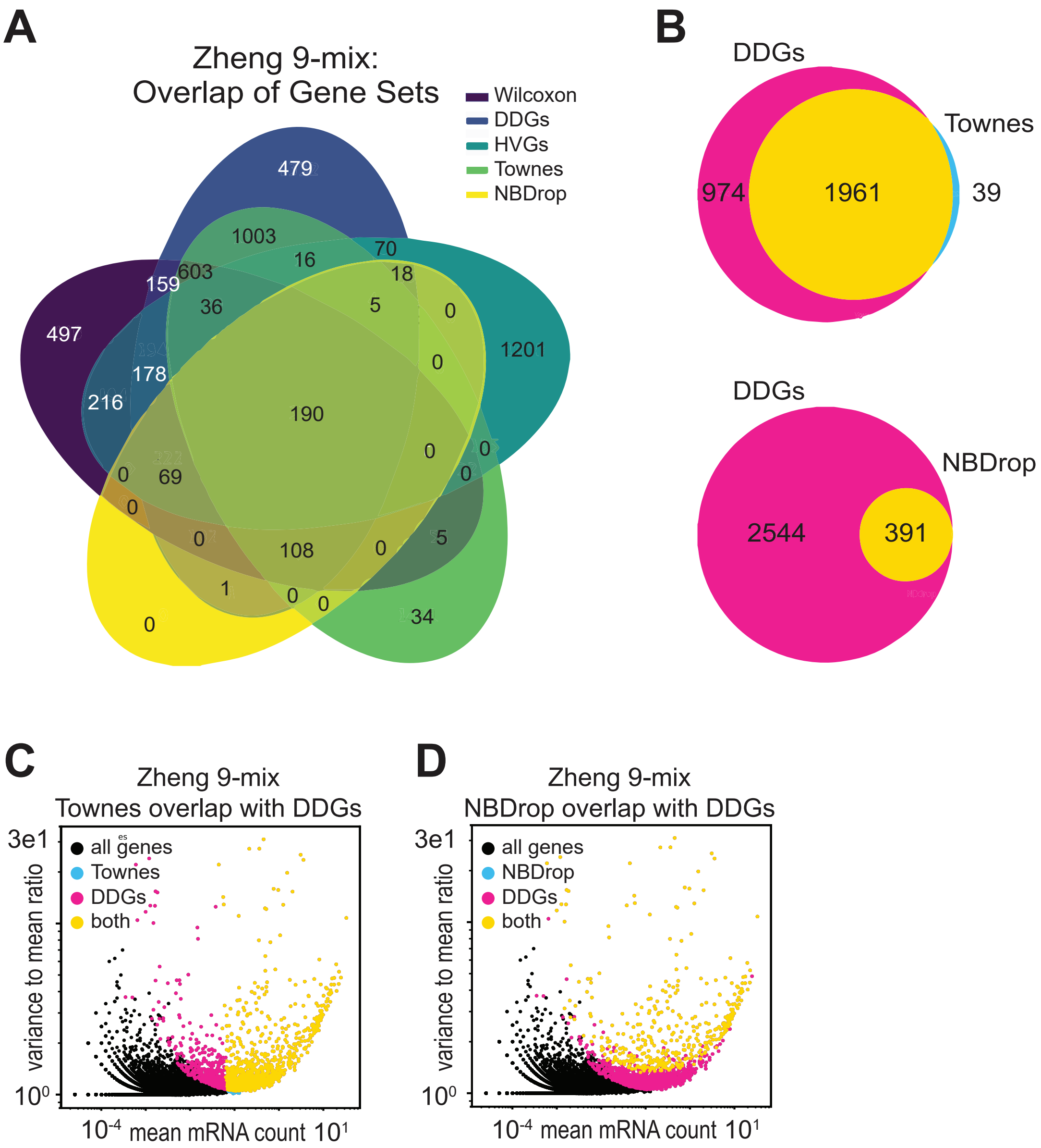


**Figure S6: A)** Venn diagram of set overlap for all feature sets calculated for the Zheng-9 lymphocyte mix. **B)** Venn diagrams of set overlap for the DDGs and the two other binomial based feature selection methods applied to the lymphocyte data. **C,D)** Scatter of the mean mRNA count versus dispersion per gene, colored by set membership for feature sets in the lymphocyte data. C) compares the Townes genes and DDGs while D) compares the NBDrop genes and DDGs.

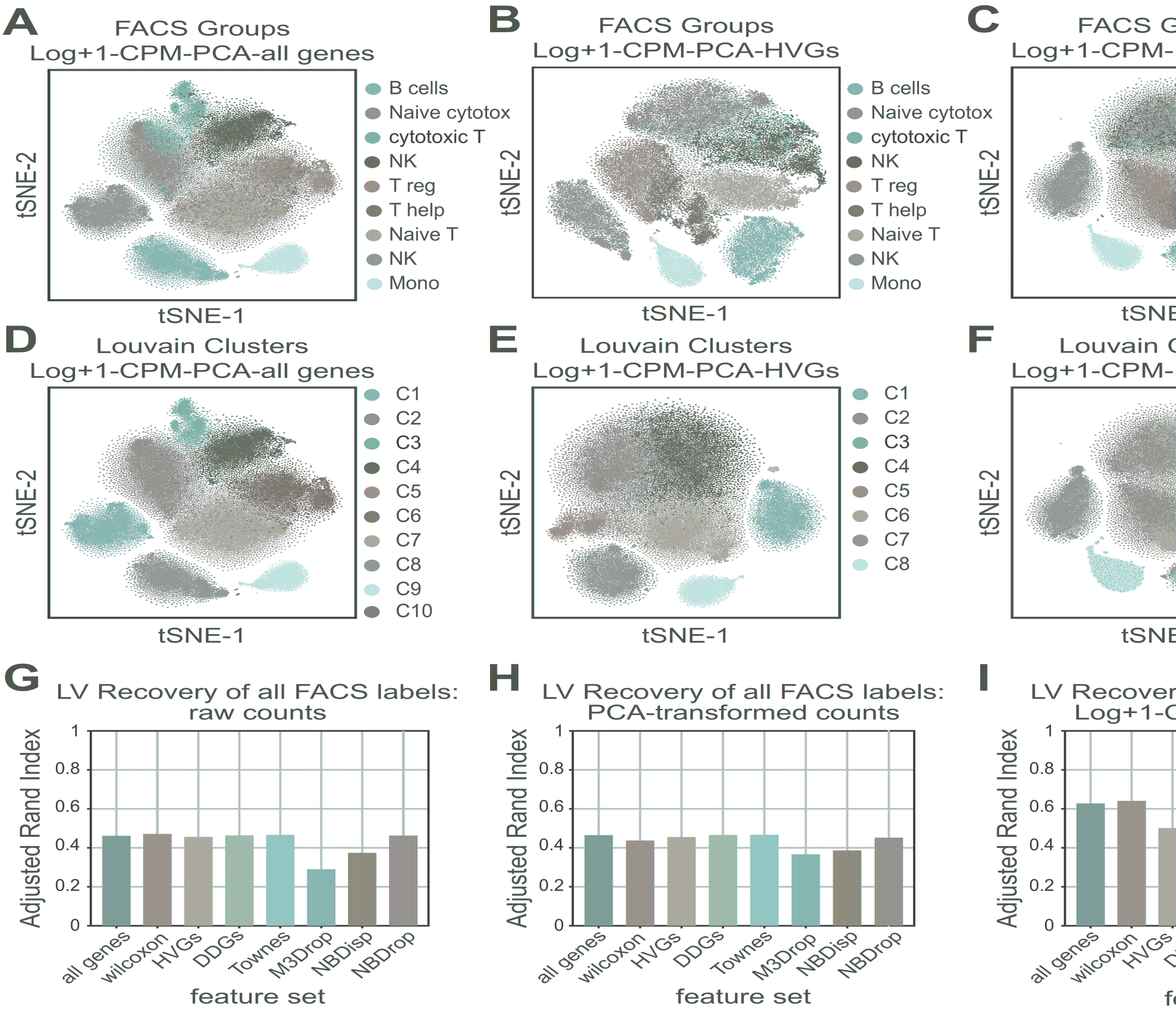


**Figure S7: A-F)** tSNE projections for dimensionality-reduced data from the Zheng-9 lymphocyte component analysis was used to reduce (A,D) all genes, (B,E) HVGs, or (C,F) DDGs. Cells are e original FACs-based set membership (A-C), or Louvain cluster membership (D-F). **G-F)** Quantific recovery by various methods of dimensionality reduction across different feature sets. Louvain cl performed across a titration of resolution parameters, cluster labels were compared to original FA highest adjusted rand index is plotted when G) raw UMI counts, H) PCA-transformed counts, or I Log+1-CPM-PCA-counts were used as a basis.

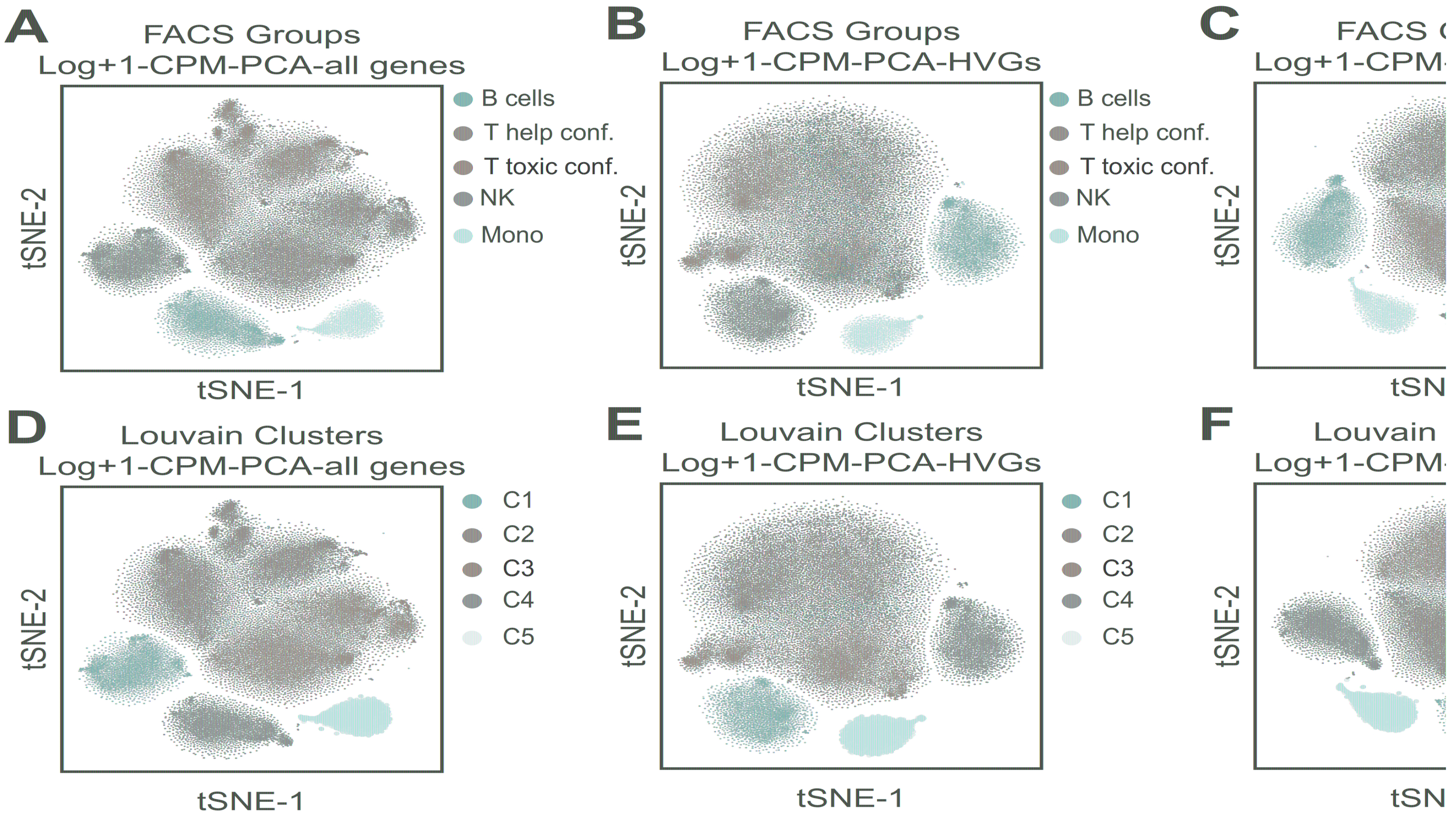


**Figure S8: A-F)** tSNE projections for dimensionality-reduced data from the Zheng-9 lymphocyt Principal component analysis was used to reduce (A,D) all genes, (B,E) HVGs, or (C,F) DDGs and CPM transforming the count data. Cells are either colored by original FACs-based set whe lineages were merged into two super sets (A-C) or Louvain cluster membership (D-F).
